# Single-cell reconstitution reveals persistence of clonal heterogeneity in the murine hematopoietic system

**DOI:** 10.1101/2021.06.28.450120

**Authors:** Nadiya Kubasova, Clara F. Alves-Pereira, Saumya Gupta, Svetlana Vinogradova, Alexander Gimelbrant, Vasco M. Barreto

**Affiliations:** CEDOC, NOVA Medical School, Faculdade de Ciências Médicas, Universidade NOVA de Lisboa, Lisboa, Portugal; Dana-Farber Cancer Institute, Department of Cancer Biology and Center of Cancer Systems Biology, Boston, MA, USA; Harvard Medical School, Department of Genetics, Boston, MA, USA; Broad Institute of MIT and Harvard, Cambridge, MA, USA; School of Genetics and Microbiology, Trinity College Dublin, Dublin

**Author notes:** Co-corresponding authors, Correspondence: Alexander Gimelbrant and Vasco M. Barreto. These authors contributed equally to this work (Nadiya Kubasova, Clara F. Alves-Pereira).

## Abstract

The persistence of patterns of monoallelic expression is a controversial matter. We report a genome-wide *in vivo* transcriptomics approach based on allelic expression imbalance to evaluate whether the transcriptional allelic patterns of single murine hematopoietic stem cells (HSC) are still present in the respective differentiated clonal B-cell populations. For 14 genes, we show conclusive evidence for a remarkable persistence in HSC-derived B clonal cells of allele-specific autosomal transcriptional states already present in HSCs. In a striking contrast to the frequency of genes with clonal allelic expression differences in clones expanded without differentiation (up to 10%), we find that clones that have undergone multiple differentiation steps *in vivo* are more similar to each other. These data suggest that most of the random allele-specific stable transcriptional states on autosomal chromosomes are established *de novo* during cell lineage differentiation. Given that allele-specific transcriptional states are more stable in cells not undergoing extensive differentiation than in the clones we assessed after full lineage differentiation *in vivo*, we introduce the “*Punctuated Disequilibria”* model: random allelic expression biases are stable if the cells are not undergoing differentiation, but may change during differentiation between developmental stages and reach a new stable equilibrium that will only be challenged if the cell engages in further differentiation. Thus, the transcriptional allelic states may not be a stable feature of the differentiating clone, but phenotypic diversity between clones of a population at any given stage of the cell lineage is still ensured.

## INTRODUCTION

One of the most remarkable features of multicellular organisms is the diversity of cellular phenotypes within each body. Isogenic cells display distinct phenotypes due to different epigenetic features or chromatin states that underlie specific gene expression programs. Technical progress in next-generation sequencing (NGS) methods has produced a wealth of data on the transcriptomics and genome-wide chromatin states of different lineages and stages within each lineage. However, distinguishing stable and reversible modes of gene regulation remains a challenge ^1^. Likewise, the epigenetic and functional inter-clonal diversity within cell lineages has been difficult to capture. One proxy for approaching these questions is to explore the allelic differences in expression.

Diploid eukaryotic organisms inherit one allele from each parent and, in most cases, the two alleles of each gene are expressed at the same time and roughly similar levels in each cell. Exceptions to this biallelic expression pattern arise from asymmetries between the two alleles, leading to unequal expression of two alleles, which can be quantified as allelic imbalances (AI). AI can have a genetic basis, due to inherited differences in each allele’s cis-regulatory regions or acquired somatic DNA modifications or be caused by allele-specific epigenetic differences accumulated by the somatic cell. Parent-of-origin genomic imprinting ^2^ and X-chromosome inactivation (XCI) ^3^, the most well-studied examples of AI due to epigenetic differences, cannot shed light on inter-clonal lineage diversity; in the former process, all somatic cells from the organism are virtually identical concerning the genomic imprint; in the latter, only two different cell populations emerge in females (differing in which X chromosome was inactivated). Potentially more useful are the random epigenetic-based AI that have been identified in autosomal genes at frequencies ranging from 2% per cell type to up to 15% of all genes ^4–8^. Some cells may express mostly or exclusively (monoallelically) one allele of these autosomal genes, whereas other cells express mostly or exclusively the other allele, a phenomenon known as random monoallelic expression (RME). These imbalances in heterozygous organisms establish clones within each lineage with phenotypic and functional differences, as in the extensively studied antigen and olfactory receptor gene ^9,10^. However, it remains to be addressed if the concept applies broadly at the functional level to more genes ^4^ and what is the real potential for clonal diversity based on the combinations of genes with distinct allelic expression levels.

Most of the studies reporting measurable frequencies of autosomal genes with random AI were performed in collections of clones expanded *in vitro*. In most cases these clones were expanded without undergoing differentiation or under limited differentiation. Building upon previous work ^11^, here we report the first genome-wide analysis of B and T cell populations emerging *in vivo* from a single hematopoietic stem cell (HSC) to evaluate whether regions in the autosomal chromosomes can keep stable expression patterns after extensive differentiation.

## RESULTS

### A single HSC with long-term reconstitution gives rise to myeloid and lymphoid cells in the blood

This work’s main goal is to study stable transcriptional states using the allelic transcriptional states of readouts in a clonal system recreated *in vivo*. For this purpose, we introduced single HSCs from a donor female mouse carrying the Ly5.2 pan-leukocyte marker in a sub-lethally irradiated recipient female mouse carrying the Ly5.1 marker to distinguish recipient and donor cells (**Supplementary Fig. 1**). The donor female F1 mice obtained by crossing B6 females with CAST males are characterized by high heterozygosity - high single nucleotide polymorphisms (SNPs) density - across the genome ^12^: about 1 SNP per 80 bp of non-repetitive genome sequence, on average, therefore enabling allele-specific analyses. The transplanted cell was left to expand and differentiate *in vivo*, producing clonal multilineage cell populations derived from a single HSC. In parallel, 50 or 200 HSCs were also transplanted per animal to generate oligoclonal or polyclonal control populations (**Fig. 1A**).

**Figure 1.**
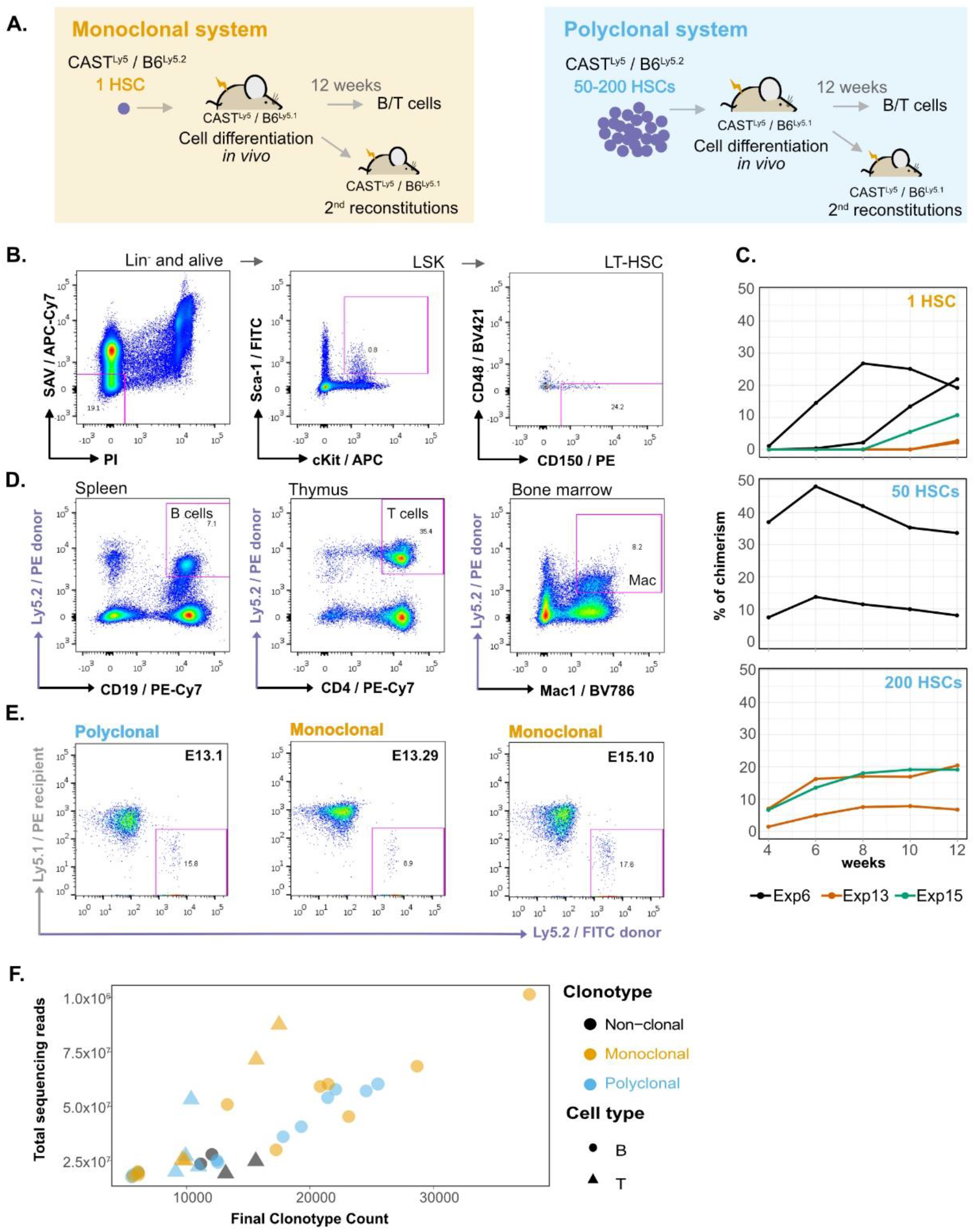
A single HSC gives rise to myeloid and lymphoid cells in the blood with long-term reconstitution. **(A)** Establishment of monoclonal and polyclonal hematopoietic systems *in vivo*. A single hematopoietic stem cell (HSC) or 50–200 hematopoietic stem cells (HSCs) were injected in sub-lethally irradiated recipient mice to generate a monoclonal or a polyclonal hematopoietic system, respectively. Both donor and recipient animals were the F1 progeny of CAST × B6 crosses, but the recipient and donor cells could be distinguished by the presence of a polymorphism in the pan-leukocyte antigen Ly5 [donor animals: F1(CAST^Ly5/Ly5^xB6^Ly5.2/Ly5.2^), recipient animals: F1(CAST^Ly5/Ly5^xB6^Ly5.2/Ly5.1^)]. Secondary reconstitutions and isolation of B/T cell populations were performed after 12 weeks of cell differentiation *in vivo*. **(B)** Long-term Hematopoietic Stem Cell (LT-HSC) isolation. The bone marrow cells of an F1 CAST^Ly5/Ly5^xB6^Ly5.2/Ly5.2^ mouse were stained with a cocktail of biotin-conjugated antibodies for surface markers of lineage-committed cells (anti-B220, anti-CD19, anti-Mac1, anti-Ter119, anti-Gr1, and anti-CD3), and subsequently, lineage-marked cells were depleted using MACS Streptavidin MicroBeads. After depletion, cells were stained with fluorophore-conjugated antibodies: APC-conjugated anti-c-Kit, FITC-conjugated anti-Sca-1, BV421-conjugated anti-CD48, PE-conjugated anti-CD150, Streptavidin-APC-Cy7, and PI, and sorted on a FACSAria. The cells were gated for PI^−^ / APC-Cy7- to exclude dead cells and any remaining lineage-positive cells, then for c-Kit^+^/Sca-1^+^ to obtain Lin-Sca^+^cKit^+^ (LSK) cells, and finally gated for CD48-/CD150^+^ to obtain LT-HSCs^14^. **(C)** Evolution of donor-derived cell populations percentages over time in the peripheral blood of the recipient animals. After blood collection, red cells were lysed, stained for Ly5.2 cells, and analyzed in a FACSCanto or FACScan instrument. **(D)** A single donor HSC differentiates into lymphoid and myeloid hematopoietic populations *in vivo*. Cells from different hematopoietic organs of recipient animals were isolated, stained, and gated on PI^−^, FITC anti-Ly5.1^+^, PE anti-Ly5.2- and PE-Cy7 anti-CD19^+^ (spleen), PE-Cy7 anti-CD4^+^ (thymus), or BV786 anti-Mac1^+^ (bone marrow). **(E)** A single donor HSC repopulates secondary recipients. Representative plots of secondary reconstitutions four weeks post-reconstitution with bone marrow cells isolated from polyclonal and monoclonal primary reconstituted animals. Blood samples of secondary reconstituted mice were lysed for red cells, stained with FITC-conjugated anti-Ly5.2 for donor cells, and PE-conjugated anti-Ly5.1 for recipient cells and analyzed using FACSCanto. **(F)** VDJ clonotypes in HSC samples. VDJ rearrangements were plotted against sequenced reads to compare the number of clonotypes in different clonal sample types.

The HSC population is heterogeneous, and several protocols based on flow cytometry were developed to distinguish between long-term HSCs (LT-HSCs) and short-term HSCs (ST-HSCs) ^13^. We used CD150^+^ and CD48^−^signaling lymphocyte activation molecule family markers on lineage negative and Sca-1^+^/cKit^+^ (LSK) cells isolated from the bone marrow of donor mouse ^14^ to single sort the LT-HSC population (**Fig. 1B**). Pure single HSCs were introduced by intravenous retro-orbital injection into recipient mice. The presence of donor cells was evaluated over 12 weeks by identifying the Ly5.2^+^ cells in the blood of recipient mice (**Fig. 1C**). From 16 experiments, 12 weeks after injections, we were able to reconstitute with a single HSC 6.6% of recipient mice with a percentage of blood chimerism in the 1– 44% range, whereas for mice injected with 50 or 200 HSCs, on average 72.9% were reconstituted and the blood chimerism was in the 2.2–87.7% range (**Supplementary Fig. 2 and Supplementary Table 1**).

Twelve weeks after injection, the animals with chimerism were sacrificed to isolate HSC derived splenic donor B cells (CD19^+^IgM^+^), donor thymocytes (CD4^+^CD8^+^), and myeloid cell populations from monoclonal and polyclonal animals (**Supplementary Fig. 3 and Fig. 1D**). We used bone marrow cells to produce secondary reconstitutions (**Fig. 1E**), showing that these CD150^+^/CD48^−^HSCs originate long-term and multilineage reconstitutions. RNA isolation and whole transcriptome sequencing were performed for the HSC derived B and T cell samples from the reconstituted animals and B cells and T cells from an unmanipulated donor female, which were used as additional non-clonal controls.

To compare the populations of evolving lymphocytes in the single-HSC and control reconstituted animals, we used MiXCR-3.0.12 ^15,16^ to quantify the V(D)J rearrangement clonotypes of sorted B and T cell samples. We observed roughly the same number of rearrangements in the single-HSC reconstitution samples, the samples produced from 50–200 HSCs, and the non-clonal samples, suggesting that there is a substantial cellular expansion in the single-HSC derived hematopoietic system before V(D)J rearrangement, which first occurs in pro-B and pro-T cells (**Fig. 1F**).

### Single HSC reconstitutions produce clonal hematopoietic systems

HSCs isolated from one donor mouse (F1[CAST^Ly5/Ly5^ × B6^Ly5.2/Ly5.2^]) were injected in multiple recipient animals (F1[CAST^Ly5/Ly5^ × B6^Ly5.2/Ly5.1^]), and allowed to expand *in vivo*. HSC-derived B cells from polyclonal and monoclonal animals for three different experiments (E6, E13, and E15) were FACS-sorted and cDNA was sequenced (RNA-Seq); for experiment 13, HSC-derived T cells were also sorted and sequenced. B and T cells from one unmanipulated donor animal were used as non-clonal control populations (**Fig. 2A**). We took advantage of XCI to internally confirm the monoclonality vs. oligo or polyclonality of the reconstitutions. A single HSC produced not only multilineage long-term reconstitutions but also hematopoietic cell populations that are clonal. In a hematopoietic system derived from a single female HSC, all cells must have inactivated the same X-chromosome, producing a complete skewing of the maternal and paternal X-linked AI (maternal allele/(maternal + paternal alleles)), which will be equal to 1 or 0; AI will tend to 0.5 as the number of clones increases. Given that the *Xist* non-coding RNA is only expressed from the inactivated X, we first performed Sanger sequencing on *Xist* cDNA, focusing on two strain-specific SNPs. As expected, chromatograms show two overlapping peaks for the control animals, whereas, only one peak was observed in the chromatogram of single-HSC reconstituted animals (**Supplementary Fig. 4**). We then deepened this analysis by calculating the AI for the X-linked genes from the NGS transcriptomics data. As expected, in the control animals, the AI values are not extreme and in some samples they are fairly balanced (close to 0.5), whereas in the single-HSC derived hematopoietic system mice the AI for the vast majority of the X-linked genes is extreme (**Fig. 2B**). Intriguingly, in samples from some single-HSC reconstituted animals, notably E13.24_B and E13.29_B, the median AI value is slightly below one. Three scenarios were considered to explain this puzzling observation: 1) more than one HSC may have erroneously be injected in these mice; 2) XCI could be leaky in the sorted lymphocytes, given that inactivated X of mature naïve T and B cells has been reported to lack the typical heterochromatic modifications ^17^; 3) contaminating recipient (polyclonal) cells were present in the sorting cells. To sort out these hypotheses, we quantified the Ly5.1 and Ly5.2 SNPs in the NGS data. Half of the samples (n=8) had around 1% of contaminating recipient cells; two samples had contaminating cells in the 2.5– 5% range, and E13.24_B and E13.29_B had contaminating cells in the 5–10% range (**Supplementary Fig. 5**). Since E13.24_B and E13.29_B are precisely the samples with the most noticeable median AI deviation from 1, we conclude that the injections were indeed with single HSCs and that the data do not support the hypothesis that XCI in lymphocytes is leaky. Thus, the dataset is composed of monoclonal samples with a low frequency of contaminating cells and oligoclonal or polyclonal control samples.

**Figure 2.**
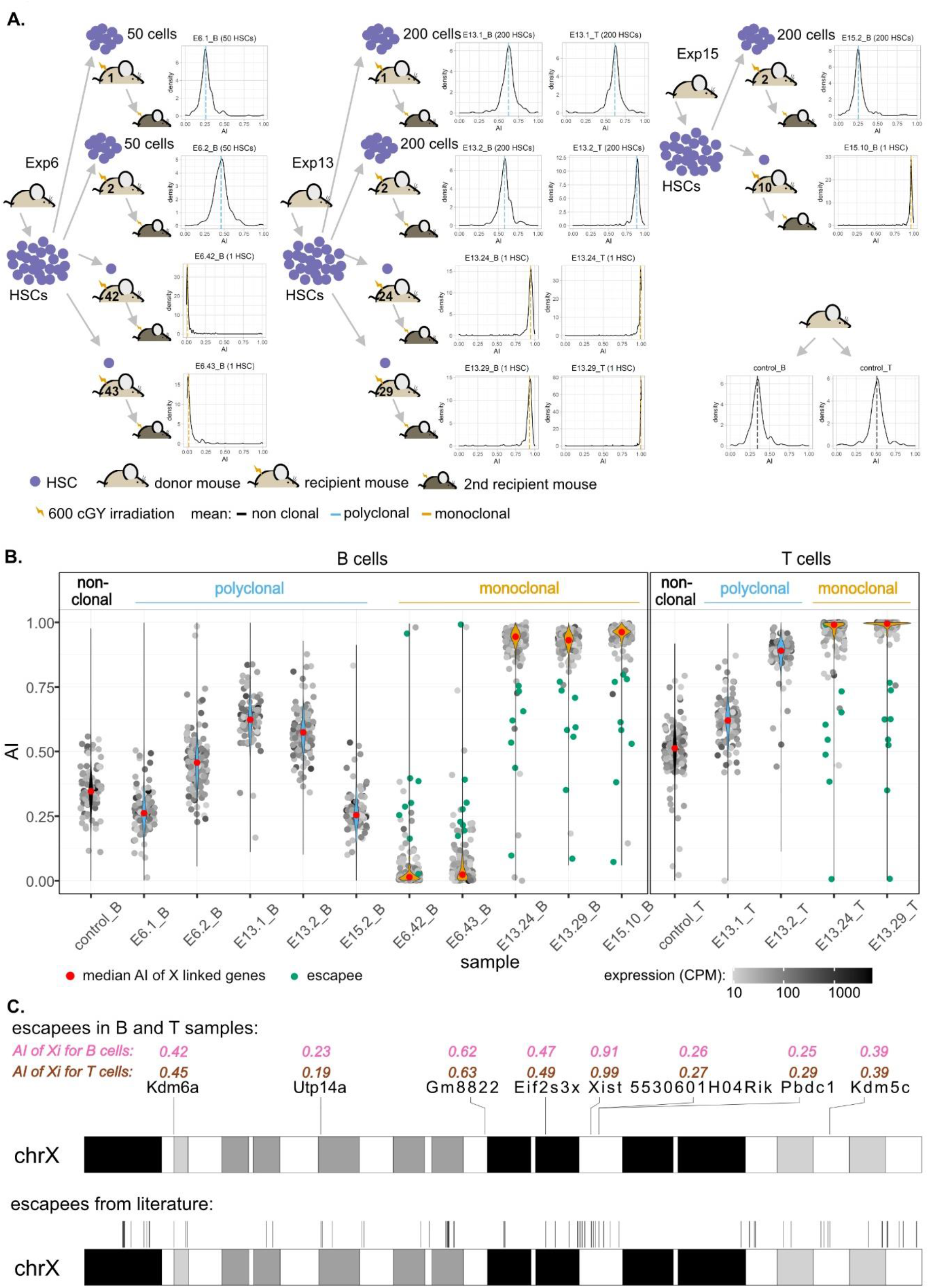
Single HSCs reconstitutions produce clonal hematopoietic systems. **(A)** Schematic representation of single and multiple HSC reconstitutions that originated the samples used for RNA-sequencing in this study (experiments E6, E13, and E15). In each experiment, HSC cells isolated from one donor mouse F1(CAST^Ly5/Ly5^xB6^Ly5.2/Ly5.2^) were injected in multiple recipient animals F1(CAST^Ly5/Ly5^xB6^Ly5.2/Ly5.1^). All animals showed long-term reconstitutions, and both monoclonal and polyclonal cells from primary repopulated animals reconstituted a secondary recipient (see representative cytometry profiles in Figure 1). The density plots represent the allelic ratios of X-chromosome linked genes for each sample, as measured by RNA-Seq. **(B)** AI of X-linked genes and XCI escapees. Violin plots superimposing dot plots of X linked genes allelic ratios per clonal/polyclonal sample. For grey dots, the opacity reflects the relative abundance in allelic counts. Genes significantly escaping X Chromosome Inactivation (XCI) (green dots) were identified by comparing the allelic ratio of that gene with a sample-corrected threshold and applying the binomial test with QCC correction ^30^. Briefly, the AI per each gene (measured by the ratio of maternal allele counts / [maternal counts + paternal counts]) were compared with a threshold value, calculated per sample, as 10% of the expression from the inactivated X chromosome (determined by the skewing observed in the entire X linked population) + the median of AI for the distributions of all the X linked genes in the sample. **(C)** Ideogram of XCI escapee genes on B and T cells are plotted along the X chromosome ideogram.

### Murine X-linked escapees identified by single-HSC reconstitutions

Genes expressed from both the active and inactive X chromosomes are known as XCI escapees. In mice, XCI escapees have been studied using three systems: 1) single-cell RNA-seq ^18,19^; 2) heterozygous female mice knockout for X-linked genes, such as *Xist* or *Hprt* ^20,21^ or heterozygous female mice for an X-linked gene linked to a reporter ^22^; 3) and clonal female F1 hybrid cell lines ^23–25^. We sought to determine whether single-HSC reconstitution could be an additional strategy to identify hematopoietic lineage-specific X escapees. X-linked genes with expression from the Xi of at least 10% of total expression ^26^ were identified taking into account the recipient cell contamination in each monoclonal sample (**Fig. 2B;** see Methods). We identified a total of eight escapees, which were escapees both in B and T samples: *5530601H04Rik*, *Eif2s3x*, *Gm8822*, *Kdm5c*, *Kdm6a*, *Pbdc1*, *Utp14a,* and *Xist* (**Supplementary Fig. 6**). These genes were plotted along the X chromosome and, as verified before ^20^, they are not clustered (**Fig. 2C**). Considering the literature, 117 genes have been described as XCI-escapees in different mouse tissues and cell lines ^20–23^. Some of these genes were excluded from our analysis for lack of expression (36 genes), insufficient number of SNPs to estimate AI (2 genes), or for not being listed in the annotation reference used in this work (1 gene). Overall, 79 genes known to escape XCI were considered. 7 of the escapees identified in our B and T samples are in this group of 79 genes; the only exception is *Gm8822*, which we have identified as an XCI pseudogene escapee and was not the subject of investigation in other studies. According to our analysis, 71 of the known escapees are not escapees in lymphocytes, which is consistent with the notion of tissue-specific XCI (**Supplementary Table 2**). Overall, we show that single-HSC transfer is an effective method to study lineage-specific XCI in blood cells.

### The vast majority of mitotically stable allelic biases of the hematopoietic system are not established during the HSC stage

To test the genome for the presence of autosomal regions in B and T cells with stable monoallelic patterns of expression reminiscent of Xi (able to persist even after an extensive program of differentiation), we generated pairwise AI comparisons of monoclonal vs. polyclonal samples, polyclonal vs. polyclonal samples; and monoclonal vs. monoclonal samples (**Fig. 3A and Supplementary Fig. 7)**. A comparison of identical samples should align all genes over the diagonal; deviations from the diagonal indicate differences in AI between the samples for a given gene. For each comparison, a Pearson’s coefficient correlation of AI for all pairwise comparisons between samples, as well as the number of genes with a significant differential AI in each pairwise comparison after applying QCC correction on the binomial test were calculated (**Fig. 3B**). If the samples from the monoclonal mice kept epigenetic states in autosomal regions in a clone-specific manner, then the correlations involving at least one monoclonal sample would be lower than the correlations found for the comparisons between controls. This was not observed. Likewise, analysis by t-distributed stochastic neighbor embedding (t-SNE) ^27^, an algorithm for visualization of high-dimensional data in a low-dimensional space, of the AI for autosomal genes would have revealed a cluster of control samples and, if each clonal line kept distinct epigenetic states, the monoclonal samples would display a more scattered distribution (**Fig. 3C)**. Again, this was not observed. We conclude that the regions in the autosomal chromosomes behaving like the X chromosomes in terms of the stable transcriptional states may not exist or represent only a small proportion of the genome that cannot be detected using this analysis.

**Figure 3.**
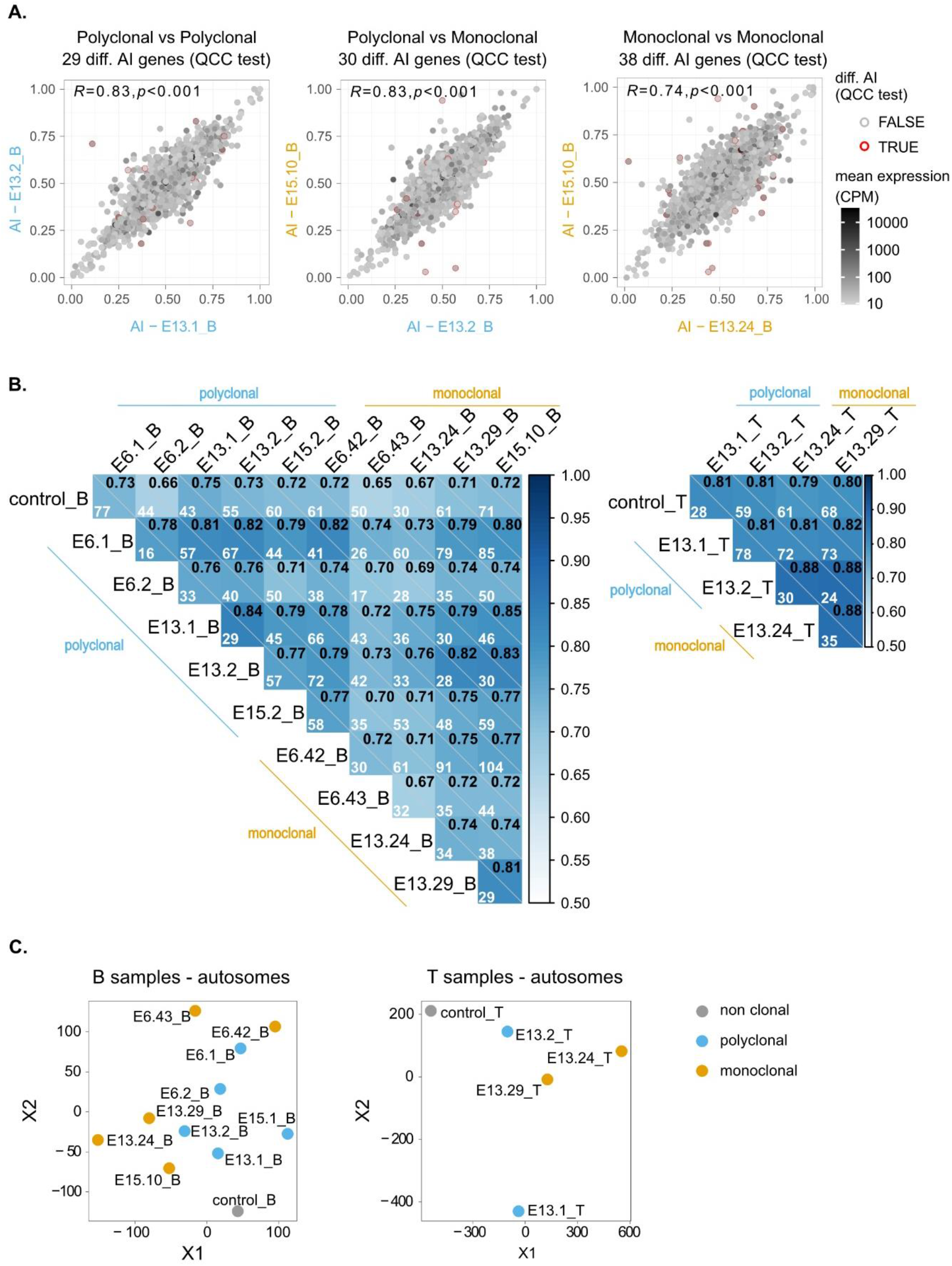
The vast majority of mitotically stable allelic biases of the hematopoietic system are not established during the HSC stage. **(A)** Representative plots of pairwise comparisons of AI between monoclonal vs. polyclonal samples, polyclonal vs. polyclonal samples; and monoclonal vs. monoclonal samples. Red circles signal the genes for which differential AI remained statistically significant after QCC correction, and the total number of these genes per comparison is shown above each plot. The Pearson’s coefficient correlation for all AI pairwise comparisons are also shown, at the upper left corner of each dot plot. A greyscale coloring the dots represents mean expression between the two samples, calculated from each sample’s CPM. **(B)** Correlograms for B and T samples. Pearson’s coefficient correlation of AI for all pairwise comparisons between samples. Within each square, Pearson’s coefficient is represented in the upper right corner, and the number of genes with a significant differential AI in each pairwise comparison after applying QCC correction on the binomial test is also shown. **(C)** Visualization of high-dimensional data of autosomal allelic imbalance in a low-dimensional space using (t-SNE algorithm) fails to show major differences between the dispersion of the polyclonal and monoclonal subsets.

### Stable transcriptional states of HSC-origin persist in the differentiated B cells for a small number of genes

The previous analysis would fail to detect a small percentage of genes with stable epigenetic states. If a gene has clone-specific AI, then the dispersion of the AI values in monoclonal samples would be higher than in the control group. To further scrutinize the dataset, we plotted the AI standard deviations of B-cell monoclonal (x-axis) and polyclonal (y-axis) samples. The plot highlighted 14 genes with higher dispersion values in the monoclonal set than in the polyclonal set (**Fig. 4A**). The fact that, above a threshold of standard deviation, no gene is found to have a standard deviation in the polyclonal set remarkably higher than in the monoclonal set suggests that the identified genes are not exceptions due to the multiple comparisons that were performed (p < 2.7 × 10^−6^, one-sided Wilcoxon test). The representation of these genes’ AI values for each animal confirms the higher dispersion in the monoclonal group compared to the polyclonal group (**Fig. 4B**). However, before these genes can be described as carrying stable epigenetic states, the possibility that these few examples result from the loss of heterozygosity (LOH) events should be addressed. In the clonal mice, during the initial stage of reconstitution, when the number of progenitor cells is low, any genetic event in a progenitor cell affecting an allele’s expression could have a sizable impact on the AI levels of the emerging populations. Thus, we performed exome sequencing in a subset of samples to evaluate whether B6 and CAST’s exons are equally represented for these 14 genes (**Fig. 4C**). The data revealed no obvious LOH for any of the genes involved. In addition, these 14 genes have not been associated with LOH or replication fragile sites and lack the molecular features typically associated with these regions, such as high expression levels and a large size ^28,29^. We conclude that the high standard deviation of the AI values for these 14 genes is not a result of LOH and is likely to reflect stable transcriptional biases originally present in the cloned HSC.

**Figure 4.**
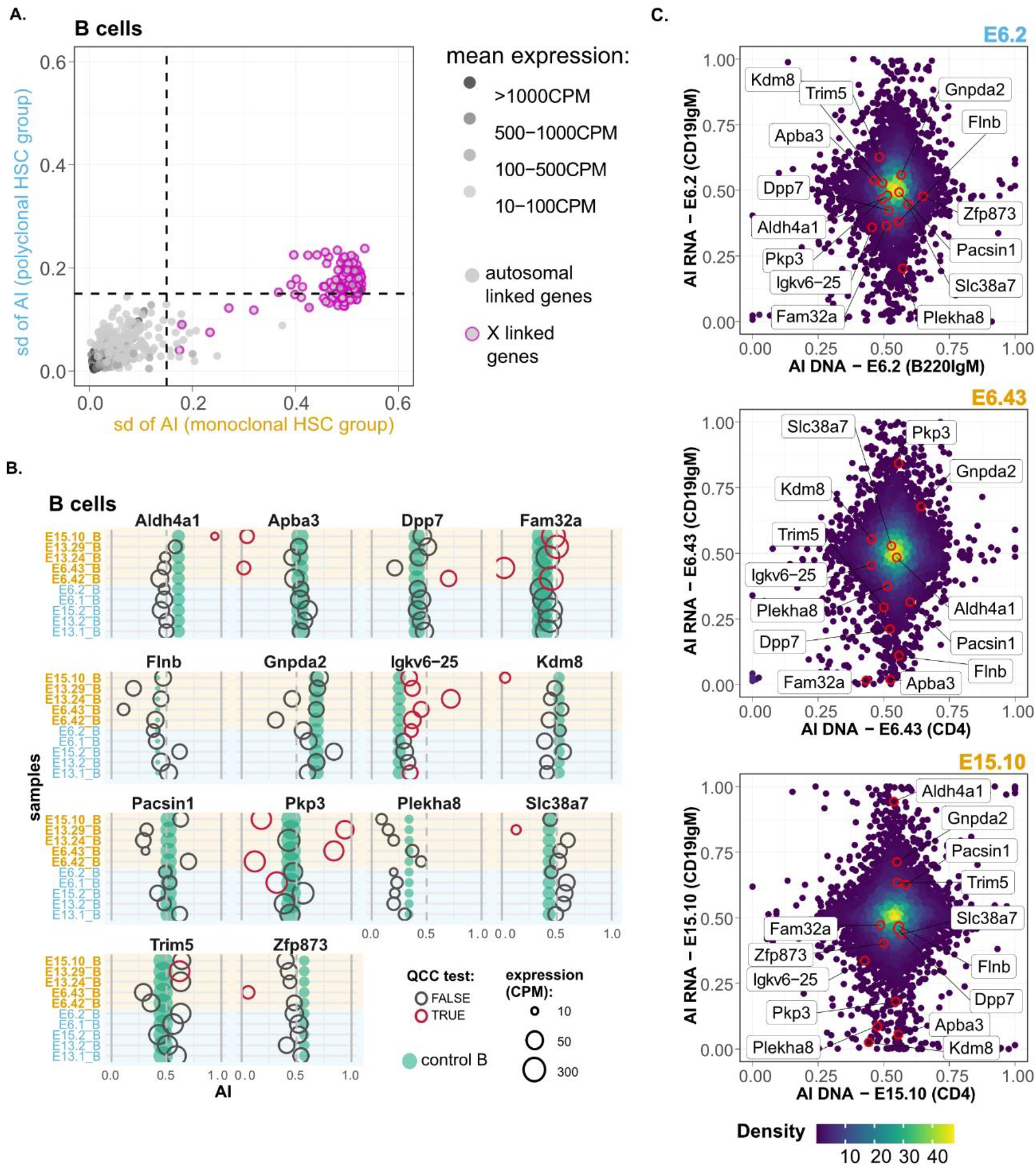
In some loci, the memory of allele-specific gene regulatory state persists over many cell divisions throughout hematopoiesis. **(A)** Dot plot showing standard deviations (SD) of AI for five B-cell monoclonal samples (x-axis) against the AI SD for five polyclonal samples (y-axis). Dashed vertical and horizontal lines - arbitrarily set at an AI SD of 0.15 - represent the threshold above which genes were considered as potentially intrinsically imbalanced. Pink-circled dots represent the X-linked genes. **(B)** Comparison of putative transcriptionally stable AI genes between all samples and non-clonal control B. Green dots are AI point estimation of control samples, and empty circles are AI point estimation of monoclonal or polyclonal samples. Red circles represent comparisons for which AI differences remained statistically significant after QCC correction. The diameter of dots/circles is proportional to expression, in CPM. **(C)** Allelic imbalance from RNA-seq data plotted against allelic imbalance from whole-exome sequencing data for the same samples (polyclonal sample E6.2 and monoclonal samples E6.43 and E15.10). Only genes with CPM>10 are represented.

### Abelson clones show a higher number of genes with clonal specific AI than lymphocytes differentiated from a single HSC

Our central question is to what extent allele-specific expression states persist in clonal populations over multiple differentiation steps. Our analysis suggest that the incidence of such stable states is much lower than was previously reported in clonal cells not undergoing differentiation ^4–8^. However, in this work we used a much more stringent statistical approach to allele-specific analysis, relying on technical replicates for RNA-seq libraries to exclude false positives ^30^.This raises the possibility that the differences could be due, at least in part, to the differences in experimental and statistical procedures compared to previous studies. To exclude this potential source of discrepancy, we applied the same analytical pipeline to RNA-seq data generated from clonal cells that grew without differentiation. We used v-Abl pro-B clonal cell lines Abl.1, Abl.2, Abl.3 and Abl.4 which were derived previously from 129S1/SvImJ × Cast/EiJ F1 female mice ^5^, with two replicate RNA-seq libraries prepared and sequenced per sample. We found that all pairwise comparisons have at least fourfold more genes with significant differences (**Fig. 5A**) than the pairwise comparison of CAST/EiJ × C57BL/6 HSC-derived clones with the highest number of genes with significant differences (**Fig. 3B**). Furthermore, the AI values in the collection of Abelson clones also have a higher dispersion than the collection of the HSC-derived clones (**Fig. 5B**). It is unlikely that these massive differences result from genetic differences between 129S1 and C57BL/6 because the two strains share an ancestor after the split from CAST/EiJ ^31^. The data suggest that in clones undergoing differentiation there is erasure and intraclonal reestablishment of AI.

**Figure 5.**
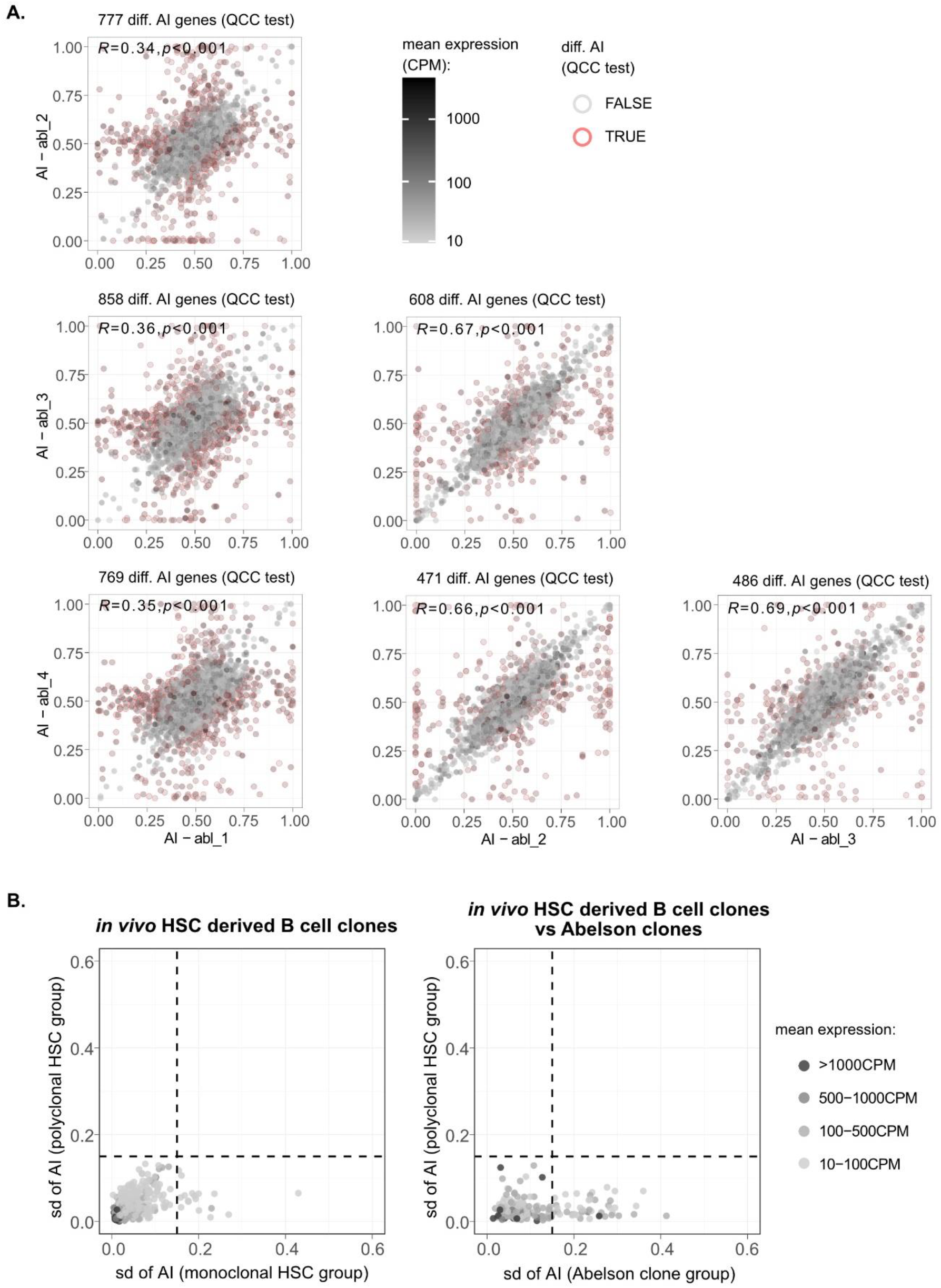
Abelson clones show a higher number of genes with clonal-specific AI than lymphocytes differentiated from a single HSC. **(A)** Representative plots of pairwise comparison of AI between different Abelson B cell clones. The Pearson’s coefficient correlation of AI and the number of genes with a significant differential AI for all pairwise comparisons between samples are shown. **(B)** Two dot plots showing standard deviations (SD) of AI for 4 monoclonal HSC-derived B cell samples (x-axis) against the AI SD for 4 polyclonal HSC-derived B cell samples (y-axis); and AI SD for all 4 Abelson clones (x-axis) against the AI SD for 4 polyclonal HSC-derived B cell samples (y-axis). Dashed vertical and horizontal lines represent the threshold above which genes were considered as potentially intrinsically imbalanced and were arbitrarily set at an AI SD of 0.15. Mean expression levels in represented as binned greyscale colors.

## DISCUSSION

There is an ongoing debate on whether phenotypic diversity due to epigenetics or somatic DNA recombination is a general phenomenon that improves the function of defined cellular populations. There is also an open discussion on the quantification of clonal RME in autosomal genes and whether this is a widespread phenomenon *in vivo* or a characteristic of clones grown *in vitro* ^32–34^. To address the latter question, we have performed a thorough analysis of random allelic expression biases in clonal B and T cell populations emerging *in vivo* after prolonged and extensive lineage differentiation in mice injected with single murine HSCs. We report two major findings. First, the analysis of these monoclonal and genetically unmanipulated hematopoietic systems allowed us to conclude that after prolonged (more than four months between HSC transfer and collection) and extensive cell division and lineage differentiation, the percentage of autosomal genes displaying RME is much lower than the estimates from collections of clones grown *in vitro* (<0.2% vs. ~2–15% ^4–8^). Second, to our knowledge, we have identified for the first time rare regions in the autosomal chromosomes that keep stable allelic transcriptional states along HSC differentiation stages. Below we discuss the implications of the technique we used and the findings for XCI, hematology, RME, and phenotypic diversity.

### XCI in a monoclonal hematopoietic system

XCI has relied on the analysis of rodent/human somatic cell hybrids ^35^, primary human cell lines ^36^, murine or human embryonic stem cells ^37,38^, murine and human-induced pluripotent stem cells ^39^, and transgenic mice with genetically engineered *Xist* locus ^21^. The former are *in vitro* systems, and the latter is an animal model in which the activation of one of the X chromosomes is imposed due to the deletion of *Xist*. Here we show that it is possible to study lineage-specific chromosome inactivation *in vivo* using genetically unmanipulated cells. Single-cell HSC reconstitution of mice identified escapees from XCI in B and T cells that had been previously identified in different tissues ^20–23^. Given the extraordinary differentiation of the hematopoietic cells from the HSCs, the interest in tissue-specific epigenetics ^40^, and the possibility of reactivation of X chromosome in lymphocytes ^17,41^, this system can be used to produce an atlas of lineage-specific XCI in the blood cells in mice and potentially also in human cells, if single human HSCs are shown to produce monoclonal human hematopoietic systems in reconstituted mice ^42^. This is currently a hot topic, as lymphocytes have been described to activate regions of the inactive X chromosome ^17,43^. The failure to observe an increased number of X escapees in lymphocytes is probably explained by the low percentage of biallelic expression in the X-linked genes in lymphocytes or the fact that the experiment was not designed to address this question.

### Autosomal versus XCI parallels

XCI and RME in autosomal regions have in common the stochastic component leading to expression vs. silencing. A number of parallels have been drawn between these phenomena ^44,45^; notably, at least one gene has been found to play a role in XCI and RME ^46^ and high concentrations of long interspersed nuclear element sequences, which were implicated in XCI ^47^, have been proposed to characterize *loci* involved in RME ^48^. Despite these possible common mechanistic features, our study establishes a fundamental difference: during lineage differentiation, RME lacks the stability of XCI.

### Applications of stable imprints in the autosomal regions

Identifying a few regions in the autosomal chromosomes with stable epigenetic states in the hematopoietic lineage could be explored in the future to develop clonality assays for the hematopoietic system. These assays have typically relied on finding significant skewing of the XCI ratio from the 1:1 ratio, which is limited to females and has a low resolution ^49^. By focusing on polymorphisms in the autosomal regions with stable epigenetic states, it should be possible to design clonality assays for both sexes that are more sensitive to decreases in clonality than the assays based on XCI.

### Punctuated Disequilibria

As a way to reconcile the lack of AI in extensively differentiated *in vivo* grown clones with the data on *in vitro* grown clones that do not undergo differentiation in culture, we propose that the evolutionary selection pressure shaping RME is at the level of the phenotypic diversity displayed by a cellular population, which does not absolutely require the persistence of the allelic biases at the deep memory clonal level. What should be crucial is that, within a given developmental stage, the cells forming a population keep distinct allelic biases, but these may change stochastically from one stage of differentiation to the next (within the clone as it undergoes differentiation) (**Fig. 6**). We call this model “*Punctuated Disequilibria*,” an obvious wordplay on a theory explaining the fossil record ^50^. “*Disequilibria*” refers to the existence of cells with different allelic biases within each population, whereas “*punctuated*” relates to the discrete instances along with lineage differentiation during which genes undergo changes in expression levels. We emphasize the key idea of the model: the uncoupling of population phenotypic diversity from clonal stability. These two concepts are typically seen as intertwined. For decades, the poster child examples of autosomal RME and the generation of phenotypic diversity within initially isogenic cell populations have been the antigen and odorant receptors, for which the univocal association between the phenotype and the clone or long-living cell is essential. In the case of the antigen receptors, the stability of the phenotype is required because the process of V(D)J recombination that builds a functional antigen receptor gene is coupled to stringent negative and positive cellular selection steps in the bone marrow or the thymus, and the emerging clone is not allowed to completely reinvent its antigen receptor after exiting the primary lymphoid organs. Although for a different reason, which is the preservation of the topographic map of the olfactory experience throughout life, each olfactory sensory neuron is also committed to the expression of a single odorant receptor gene (and allele). These examples of phenotypic diversity are spectacular but also exceptional in the sense that an antigen receptor gene depends on a unique process of somatic DNA recombination, and the odorant receptor genes make up the largest gene family in the mammalian genome. Less unique genes, particularly in the blood cells, which circulate permanently, may be better described within each cell population and along lineage differentiation by punctuated disequilibria rather than phenotypic clonal stability. In the future, we will address how the allelic expression equilibrium of a given gene is disturbed by time, cell cycle, the extent of differentiation, the changes in the expression levels throughout the development of the gene and its neighbor genes, and we will also dissect the epigenetic and genetic components of this process. For now, we propose that the phenotypic diversity of a given cell population could rely less on clonal stability than on the independence of each cell during the stages of gene (re)activation that punctuate lineage commitment and cell activation, which may set a new expression balance for the alleles until next stage of differentiation.

## ACKNOWLEDGMENTS

The authors would like to acknowledge Cláudia Andrade from the Facility of Flow Cytometry from CEDOC for excellent technical work, and both the Antibody Unit and the Animal House Facility from Instituto Gulbenkian de Ciência. This work was supported by the FCT (*Fundação para a Ciência e a Tecnologia*) under grants PTDC/BEX-BCM/5900/2014 and IF/01721/2014/CP1252/CT0005, and European Union’s Horizon 2020 research and innovation programme under the Marie Sklodowska-Curie grant agreement No 752806. Nadiya Kubasova received a fellowship PD/BD/114164/2016 from the FCT. We thank Ana Cumano, Anne-Valerie Gendrel, and Thiago L. Carvalho for their helpful comments.

## AUTHOR CONTRIBUTIONS

NK, CFAP, AG, and VMB designed the project. NK performed all *in vivo* experiments, prepared all figures, and wrote the methods section. SG produced the Abelson data. CFAP, NK, SV and AG analyzed the NGS data. NK, CFAP, AG, and VMB analyzed the data. VMB wrote the first draft, which was extensively edited by NK, CFAP, and AG.

## DECLARATION OF INTERESTS

The authors have no conflict of interest to disclose.

## METHODS

### Animal breeding

All mice were bred and maintained at the specific pathogen-free animal facilities of the Instituto Gulbenkian de Ciência (IGC, Oeiras, Portugal). C57BL/6J-Ly5.1 (C57BL/6J strain carrying the pan-leukocyte marker Ly5.1), C57BL/6J-Ly5.2 (C57BL/6J strain carrying the pan-leukocyte marker Ly5.2), and CAST/EiJ were originally received from The Jackson Laboratory (Bar Harbor, ME, USA). Animals used in reconstitution experiments were bred at our animal facility to generate female heterozygous F1 donor (CAST/EiJ × C57BL/6J-Ly5.2) and recipient (CAST/EiJ × C57BL/6J-Ly5.1) animals. All animals used in cell transfer experiments were 8–16 week-old. This research project was reviewed and approved by the Ethics Committee of the IGC and by the Portuguese National Entity that regulates the use of laboratory animals.

### HSCs isolation

The bone marrow was flushed out and single-cell-suspended with FACS buffer (1× PBS, 2% FBS) from the tibia and femur using a syringe. The erythrocytes were lysed with red blood cell lysis buffer (RBC lysis buffer) (155 mM NH_4_Cl, 10 mM NaHCO_3_, 0.1 mM EDTA, pH 7.3) for 5 min and immediately rinsed and washed with FACS buffer. The cells were blocked with FcBlock (anti-CD16/32) for 15 min at 4°C and washed. Enrichment for lineage negative cells was performed by incubating cell suspension with a cocktail of biotin-conjugated antibodies for surface markers of lineage-committed cells (anti-CD45R/B220, anti-CD19, anti-CD11b/Mac1, anti-Ly-76/Ter119, anti-Ly6G/Gr1, and anti-CD3) and, subsequently, lineage-marked cells were depleted using MACS Streptavidin MicroBeads (Miltenyi Biotec) for negative selection of lineage-positive cells by immunomagnetic separation using a MACS column (Miltenyi Biotec). Cells were further stained with PI and fluorophore-conjugated antibodies: APC-conjugated anti-c-Kit, PE-Cy7-conjugated anti-Sca-1, BV421-conjugated anti-CD48, PE-conjugated anti-CD150 and Streptavidin-APC-Cy7, to isolate LH-HSCs (adopted from Kiel et al. ^14^). LT-HSCs were sorted on a FACSAria II using the single-cell deposition unit into the individual wells of Terasaki plates (no. 452256, MicroWell 60-well MiniTray, Nunc Brand, Thermo Fisher Scientific Inc.) preloaded with 15 μL of FACS buffer. Each well was examined in a 4°C room using an inverted microscope and the wells with a single cell were used in the reconstitutions.

### Animal reconstitutions

8–16 week-old recipient females received sublethal whole-body g-irradiation with 600 cGy (Gammacell 2000 Mølsgaard Medical), 2–6 h before an intravenous retro-orbital injection with single-HSC or 50-200 HSCs. Recipient animals were analyzed routinely four weeks after injection and every two weeks for up to 12 weeks for the presence of chimeric cells in the peripheral blood. Blood samples were collected from the submandibular vein in EDTA, erythrocytes were lysed using RBC lysis buffer, cells were stained with PE-conjugated anti-Ly5.1 and FITC-conjugated anti-Ly5.2 antibodies, and analyzed by FACSCanto or FACScan.

### Processing of animal samples

Animals selected for subsequent analysis showed chimeric cells 12 weeks post-reconstitution were sacrificed and processed by removing thymi, spleens, and bone marrows. Single-cell suspension from bone marrow was obtained as described above using a syringe and spleen, and thymus using a 70-μm nylon mesh. Erythrocytes were lysed with RBC lysis buffer for 5 min and immediately rinsed and washed with FACS buffer. 30% of cell suspension from bone marrow was saved for reconstitution of sublethally irradiated secondary recipient female mice, injected by intravenous retro-orbital administration, and analyzed for chimerism four weeks post-injection as described above. Different stainings with labeled antibodies were used to analyze and sort lymphoid populations in the spleen and thymus and myeloid population in bone marrow or spleen with FACSAriaII, after cell blocking with FcBlock. In experiment 6, a combination of PI, APC-Cy7-conjugated anti-Ly5.1, and PE-conjugated anti-Ly5.2 was used with markers PE-Cy7-conjugated anti-CD19 APC-conjugated anti-IgM and BV786-conjugated anti-Mac1 for spleen; and PE-Cy5-conjugated anti-CD4 and BV605-conjugated anti-CD8 for thymus. In experiments 13 and 15, a combination of PI, FITC-conjugated anti-Ly5.1, and PE-conjugated anti-Ly5.2 was used with markers PE-Cy7-conjugated anti-CD19 and APC-conjugated anti-IgM for spleen; PE-Cy7-conjugated anti-CD4 and BV605-conjugated anti-CD8 for thymus, and BV786-conjugated anti-Mac1 for bone marrow.

### RNA extraction

After cell sorting, pellets were harvested by centrifugation and resuspended in 0.25 mL of TRIzol Reagent or 0.1 mL of Absolutely RNA Nanoprep Kit (Agilent #400753) lysis buffer. Homogenized samples were stored at −80°C until RNA isolation, which was performed according to the manufacturer’s protocols.

### Monoclonality screening

To test for monoclonality before sequencing, RNA was isolated from the same repopulated animals using sorted cell populations other than the sequenced ones. cDNA was prepared using SuperScript IV (ThermoFisher #18090050) following the manufacturer’s recommendations. *Xist* locus was amplified in two individual reactions using two sets of primers obtaining amplicons with two different SNPs: Fw1 5’agacgctttcctgaacccag with R1 5’aagatgctgcagtcaggc; and Fw2 5’ggagtgaagagtgctggagag with R2 5’gtcagtgccactattgcagc. PCR was performed with GoTaq DNA polymerase (Promega #M3005) using the following program: 5 min at 95°C, 45 cycles of 30 s at 95°C, 30 s at 60°C, and 25 s at 72°C, and a final elongation of 7 min at 72°C. The amplicons were separated in agarose gel, purified, and sequenced by Sanger sequencing with Fw1 or R2 primers.

### cDNA library preparation and whole-transcriptome sequencing

Omega Bioservices, USA, performed cDNA library preparation and whole transcriptome sequencing. According to the manufacturer’s protocol, RNA-sequencing libraries were prepared using SMART-Seq v4 Ultra Low Input RNA Kit (Clontech). Technical replicates of 10 ng of RNA were used as input. The RNA was primed by an oligo(dT) primer (3’ SMART-Seq CDS Primer II A), and first-strand cDNA synthesis was performed at 42°C for 90 min and 70°C for ten min. The resulting cDNA was then amplified via PCR using the following program: 1 min at 95°C, eight cycles of 10 sec at 98°C, 30 s at 65°C, and 3 min at 68°C, and a final elongation of 10 min at 72°C. 150-200 pg full-length cDNA was tagged and fragmented by the Nextera XT transposome (Illumina) and amplified by PCR: 30 s at 95 °C, 12 cycles of 10 s at 95 °C, 30 s at 55 °C, and 30 s at 72 °C, then 5 min at 72 °C. Mag-Bind RxnPure Plus magnetic beads (Omega Bio-tek) were used to purify the library and provide a size-selection step. The libraries were then pooled in equimolar concentrations and sequenced on Illumina HiSeq 2500 machine (150 bp, paired-end).

### Allele-specific gene expression analysis from RNA-Seq

RNA-Seq data analysis for AI estimation followed the ASEReadCounter* tool adapted from the GATK pipeline ^51^ for the pre-processing read alignment steps up to allele counts, and the statistical R package Qllelic.v0.3.2 for calculation of the quality control constant (QCC) and estimation of confidence intervals for differential AI analysis ^30^. RNA-seq reads were trimmed from nextera adapters with cutadapt.v.1.14 using the wrapper trim_galore. Sequencing reads were aligned to the reference genome (maternal) and imputed genome (paternal) with the STAR aligner v.2.5.4a, with default filtering parameters and accepting only uniquely aligned reads. Samtools mpileup (v.1.3.1) was used to estimate allele-specific coverage over SNPs. Gene models were generated by collapsing all exons belonging to the same gene, based on the GRCm38.68 RefSeq GTF file downloaded from ftp://ftp.ensembl.org/pub/release-68/gtf/, where overlapping regions belonging to multiple genes were excluded. Point estimates of AI for a gene were obtained as the ratio of maternal gene counts over total allelic gene counts. Gene abundance counts were obtained with featureCounts from the same bam files generated with the ASEReadCounter* alignment pipeline, and abundance was estimated with edgeR.

### XCI escapees

X-linked genes were considered XCI escapees if significant expression from the inactive X chromosome were identified in each single-HSC derived sample by comparing the allelic imbalance value with a threshold value calculated for each sample as the median of the AI distribution for all genes on that sample (to account for potential biallelic contamination) +/−0.1. The comparisons were performed by applying the binomial test with quality control correction for technical replicates (QCC) ^30^. To consider a gene as an escapee, we defined three criteria: 1) only samples with expression higher than 10 CPM (count-per-million) were considered; 2) the mean of AI in the control samples (polyclonal and non-clonal samples) was fairly balanced (0.5±0.2); 3) and AI was above the monoclonal sample threshold in at least two samples from the same tissue (B or T cells) or different in at least one B cell sample and at least one T cell sample. (**Supplementary Fig. 6**).

**Figure 6.**
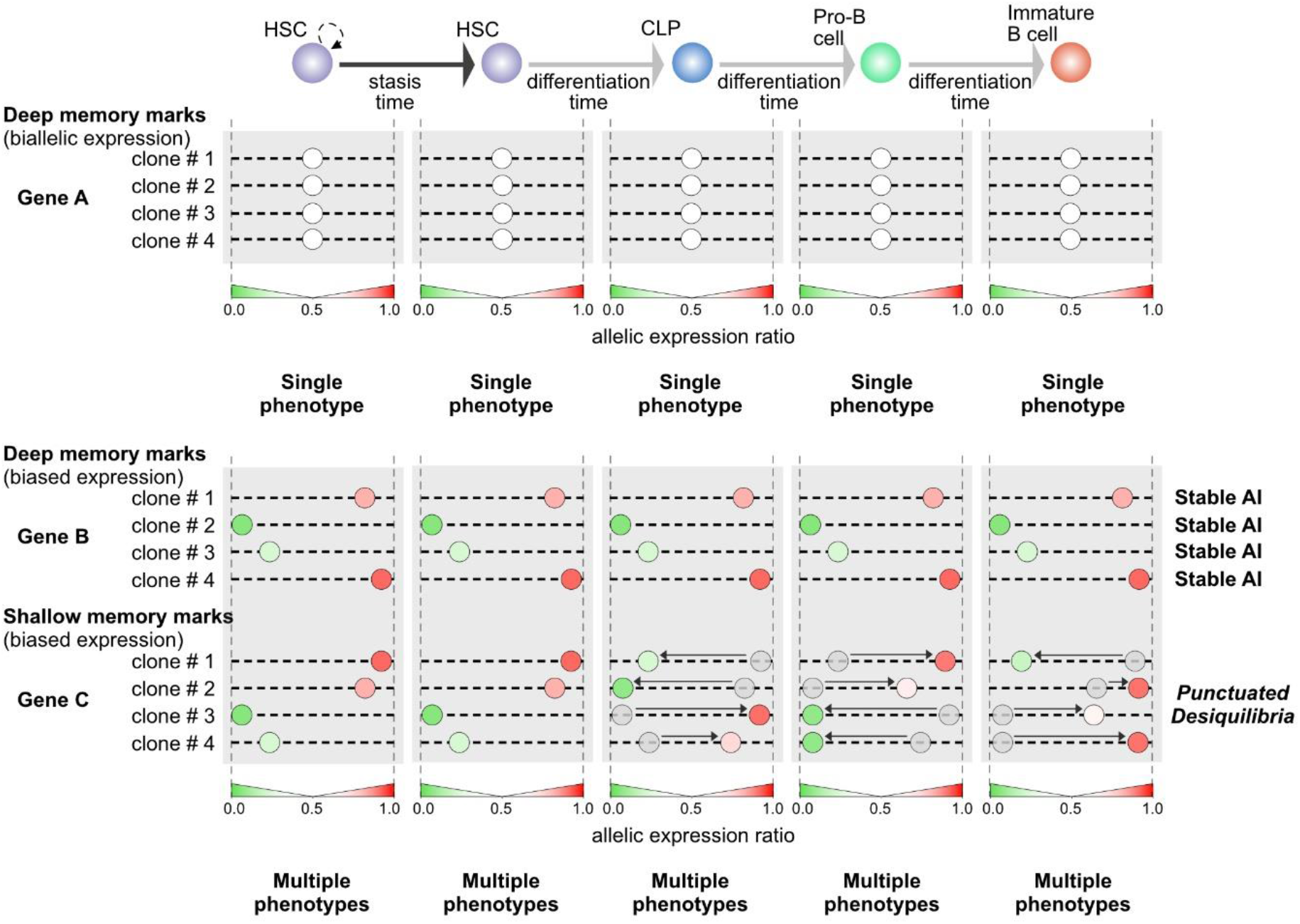
Punctuated Disequilibria model. Random allelic biases are stable if the cell and its progeny do not engage in differentiation, but may change upon differentiation and reach a new stable allelic expression equilibrium. As a result, we can subdivide mitotically stable AI according to their persistence. Shallow memory marks are stable during proliferation, but the AI values may shift during differentiation, while deep memory marksare stable even during differentiation. In our study, the analysis of B cells generated from a single HSC reveals random biases in genes with deep memory marks, but the AI biases in genes with shallow memory marks are undetected because, intraclonally, the AI values change from one cell stage to the other. The behavior of shallow marks disrupts the AI clonal stability throughout differentiation, but within each cell stage the different AI values are stable and ensure the needed single phenotype.

### VDJ clonotypes

Immunoglobulin rearrangements were detected by alignment of RNA-Seq raw data with reference germline V, D, J, and C gene sequences and assembled into clonotypes with MiXCR-3.0.12 ^15,16^.

### DNA library preparation and whole-exome sequencing

DNA was recovered from samples stored in TRIzol Reagent according to the instructions of the manufacturers, resuspended in DNase-free water, and stored at −20°C. Novogene, UK, performed DNA library preparation and whole-exome sequencing using Agilent SureSelect Mouse All ExonV6 kit (Agilent Technologies) following recommendations of manufacturer, and × index codes were added to attribute sequences to each sample. The genomic DNA samples were randomly fragmented by sonication (Covaris) to the size of 180–280 bp fragments. The remaining overhangs were converted into blunt ends via exonuclease/polymerase activities. After adenylation of 3’ ends of DNA fragments, adapter oligonucleotides were ligated. DNA fragments with ligated adapter molecules on both ends were selectively enriched in a PCR reaction. The libraries were hybridized with biotin-labeled probes, and magnetic beads with streptomycin were used to capture the exons. After washing beads and digesting the probes, the captured libraries were enriched in a PCR reaction to add index tags. The products were purified with the AMPure XP system (Beckman Coulter). DNA libraries were sequenced on an Illumina platform (150 bp, paired-end). Read alignment and allele counts were based on the ASEReadCounter* pipeline.

### Abelson clones

v-Abl pro-B clonal cell lines Abl.1, Abl.2, Abl.3 and Abl.4 were derived previously from 129S1/SvImJxCast/EiJ F1 female mice by expansion of FACS-sorted single cells after immortalization ^5^. Immortalized B-cell clonal lines were cultured in Roswell Park Memorial Institute (RPMI) medium (Gibco), containing 15% FBS (Sigma), 1X L-Glutamine (Gibco), 1X Penicillin/Streptomycin (Gibco), 0.1% β-mercaptoethanol (Sigma). Culture medium also contained 1% DMSO. On day 2 of the culture, live cells were collected after sucrose gradient centrifugation (Histopaque-1077, Sigma, Cat 10771), and RNA was extracted from cells using a magnetic bead-based protocol using Sera-Mag SpeedBeadsTM (GE Healthcare). Two libraries were prepared per clone using SMARTseqv4 kit (Clonetech), starting with 10ng input RNA for each library according to manufacturers’ instructions. Abl.1 clone was sequenced on the Illumina NextSeq 500 machine (75 bp, single-end); clones Abl.2, Abl.3 and Abl.4 were sequenced on Illumina HiSeq 4000 machine (150 bp, paired-end). RNA-seq data analysis followed the same pipeline as for HSC derived clones *in vivo*, with exception of the maternal reference genome which was 129S1. These data were originally generated for the work described in bioRxiv by Gupta et al, 2020 Preprint ^52^.

### Statistical analysis

The difference between the AI point estimates of two clones, or the difference of point estimate and a threshold (e.g., X-chr escapees), was accepted as significant after accounting for experiment-specific overdispersion of 2 replicates using the R package Qllelic.v0.3.2 ^30^.

### Data sharing statement

The entire set of HSC NGS raw data (RNA-Seq and Whole Exome Sequencing) and processed counts files have been deposited to the NCBI’s Gene Expression Omnibus database with series accession number [GEO:GSE174040]. Abelson clones RNA-Seq data have been previously deposited with series accession number [GEO: GSE144007].

## REFERENCES

1. van der Veeken, J., Zhong, Y., Sharma, R., Mazutis, L., Dao, P., Pe’er, D., Leslie, C.S. and Rudensky, A.Y. (2019). Natural Genetic Variation Reveals Key Features of Epigenetic and Transcriptional Memory in Virus-Specific CD8 T Cells. Immunity 50, 1202–1217.e7.

2. Reik, W., and Walter, J. (2001). Genomic imprinting: Parental influence on the genome. Nat. Rev. Genet. 2, 21–32.

3. Disteche, C.M., and Berletch, J.B. (2015). X-chromosome inactivation and escape. J. Genet. 94, 591–599.

4. Gimelbrant, A., Hutchinson, J.N., Thompson, B.R., and Chess, A. (2007). Widespread monoallelic expression on human autosomes. Science (80-.). 318, 1136–1140.

5. Zwemer, L.M., Zak, A., Thompson, B.R., Kirby, A., Daly, M.J., Chess, A., and Gimelbrant, A.A. (2012). Autosomal monoallelic expression in the mouse. Genome Biol. 13.

6. Jeffries, A.R., Perfect, L.W., Ledderose, J., Schalkwyk, L.C., Bray, N.J., Mill, J., and Price, J., (2012). Stochastic choice of allelic expression in human neural stem cells. Stem Cells 30, 1938–1947.

7. Gendrel, A.V., Attia, M., Chen, C.J., Diabangouaya, P., Servant, N., Barillot, E., and Heard, E., (2014). Developmental dynamics and disease potential of random monoallelic gene expression. Dev. Cell 28, 366–380.

8. Eckersley-Maslin, M.A., Thybert, D., Bergmann, J.H., Marioni, J.C., Flicek, P., and Spector, D.L., (2014). Random monoallelic gene expression increases upon embryonic stem cell differentiation. Dev. Cell 28, 351–365.

9. Vettermann, C., and Schlissel, M.S., (2010). Allelic exclusion of immunoglobulin genes: Models and mechanisms. Immunol. Rev. 237, 22–42.

10. Monahan, K., and Lomvardas, S., (2015). Monoallelic Expression of Olfactory Receptors. Annu. Rev. Cell Dev. Biol. 31, 721–740.

11. Alves-Pereira, C.F., De Freitas, R., Lopes, T., Gardner, R., Marta, F., Vieira, P., and Barreto, V.M., (2014). Independent recruitment of Igh alleles in V(D)J recombination. Nat. Commun. 5.

12. Frazer, K.A., Eskin, E., Kang, H.M., Bogue, M.A., Hinds, D.A., Beilharz, E.J., Gupta, R. V., Montgomery, J., Morenzoni, M.M., Nilsen, G.B., et al. (2007). A sequence-based variation map of 8.27 million SNPs in inbred mouse strains. Nature 448, 1050–1053.

13. Mayle, A., Luo, M., Jeong, M., and Goodell, M.A., (2013). Flow cytometry analysis of murine hematopoietic stem cells. Cytom. Part A 83A, 27–37.

14. Kiel, M.J., Yilmaz, Ö.H., Iwashita, T., Yilmaz, O.H., Terhorst, C., and Morrison, S.J. (2005). SLAM family receptors distinguish hematopoietic stem and progenitor cells and reveal endothelial niches for stem cells. Cell 121, 1109–1121.

15. Bolotin, D.A., Poslavsky, S., Mitrophanov, I., Shugay, M., Mamedov, I.Z., Putintseva, E. V., and Chudakov, D.M., (2015). MiXCR: Software for comprehensive adaptive immunity profiling. Nat. Methods 12, 380–381.

16. Bolotin, D.A., Poslavsky, S., Davydov, A.N., Frenkel, F.E., Fanchi, L., Zolotareva, O.I., Hemmers, S., Putintseva, E. V, Obraztsova, A.S., Shugay, M., et al. (2017). Antigen receptor repertoire profiling from RNA-seq data HHS Public Access Author manuscript. Nat Biotechnol 35, 908–911.

17. Wang, J., Syrett, C.M., Kramer, M.C., Basu, A., Atchison, M.L., and Anguera, M.C. (2016). Unusual maintenance of X chromosome inactivation predisposes female lymphocytes for increased expression from the inactive X. Proc. Natl. Acad. Sci. U. S. A. 113, E2029–E2038.

18. Chen, G., Schell, J.P., Benitez, J.A., Petropoulos, S., Yilmaz, M., Reinius, B., Alekseenko, Z., Shi, L., Hedlund, E., Lanner, F., et al. (2016). Single-cell analyses of X Chromosome inactivation dynamics and pluripotency during differentiation. Genome Res. 26, 1342–1354.

19. Borensztein, M., Syx, L., Ancelin, K., Diabangouaya, P., Picard, C., Liu, T., Liang, J. Bin, Vassilev, I., Galupa, R., Servant, N., et al. (2017). Xist-dependent imprinted X inactivation and the early developmental consequences of its failure. Nat. Struct. Mol. Biol. 24, 226–233.

20. Yang, F., Babak, T., Shendure, J., and Disteche, C.M., (2010). Global survey of escape from X inactivation by RNA-sequencing in mouse. Genome Res. 20, 614–622.

21. Berletch, J.B., Ma, W., Yang, F., Shendure, J., Noble, W.S., Disteche, C.M., and Deng, X. (2015). Escape from X Inactivation Varies in Mouse Tissues. PLoS Genet. 11, 1–26.

22. Wu, H., Luo, J., Yu, H., Rattner, A., Mo, A., Wang, Y., Smallwood, P.M., Erlanger, B., Wheelan, S.J., and Nathans, J., (2014). Cellular Resolution Maps of X Chromosome Inactivation: Implications for Neural Development, Function, and Disease. Neuron 81, 103–119.

23. Li, S.M., Valo, Z., Wang, J., Gao, H., Bowers, C.W., and Singer-Sam, J. (2012). Transcriptome-wide survey of mouse CNS-Derived cells reveals monoallelic expression within novel gene families. PLoS One 7, 31751.

24. Splinter, E., de Wit, E., Nora, E.P., Klous, P., van de Werken, H.J.G., Zhu, Y., Kaaij, L.J.T., van Ijcken, W., Gribnau, J., Heard, E., et al. (2011). The inactive X chromosome adopts a unique three-dimensional conformation that is dependent on Xist RNA. Genes Dev. 25, 1371–1383.

25. Calabrese, J.M., Sun, W., Song, L., Mugford, J.W., Williams, L., Yee, D., Starmer, J., Mieczkowski, P., Crawford, G.E., and Magnuson, T. (2012). Site-specific silencing of regulatory elements as a mechanism of x inactivation. Cell 151, 951–963.

26. Carrel, L., and Willard, H.F. (2005). X-inactivation profile reveals extensive variability in X-linked gene expression in females. Nature 434, 400–404.

27. Hinton, G., and Roweis, S. (2003). Stochastic neighbor embedding. In Advances in Neural Information Processing Systems, pp. 833–840.

28. Helmrich, A., Stout-Weider, K., Hermann, K., Schrock, E., and Heiden, T. (2006). Common fragile sites are conserved features of human and mouse chromosomes and relate to large active genes. Genome Res. 16, 1222–1230.

29. Barlow, J.H., Faryabi, R.B., Callén, E., Wong, N., Malhowski, A., Chen, H.T., Gutierrez-Cruz, G., Sun, H.W., McKinnon, P., Wright, G., et al. (2013). Identification of early replicating fragile sites that contribute to genome instability. Cell 152, 620–632.

30. Mendelevich, A., Vinogradova, S., Gupta, S., Mironov, A., Sunyaev, S., and Gimelbrant, A. (2021). Replicate sequencing libraries are important for quantification of allelic imbalance. Nat. Commun. in press.

31. Witmer, P.D., Doheny, K.F., Adams, M.K., Boehm, C.D., Dizon, J.S., Goldstein, J.L., Templeton, T.M., Wheaton, A.M., Dong, P.N., Pugh, E.W., et al. (2003). The development of a highly informative mouse simple sequence length polymorphism (SSLP) marker set and construction of a mouse family tree using parsimony analysis. Genome Res. 13, 485–491.

32. RV, P., Sundaresh, A., Karunyaa, M., Arun, A., and Gayen, S. (2021). Autosomal Clonal Monoallelic Expression: Natural or Artifactual? Trends Genet. 37, 206–211.

33. Vigneau, S., Vinogradova, S., Savova, V., and Gimelbrant, A. (2018). High prevalence of clonal monoallelic expression. Nat. Genet. 50, 1198–1199.

34. Reinius, B., and Sandberg, R. (2018). Reply to ‘High prevalence of clonal monoallelic expression.’ Nat. Genet. 50, 1199–1200.

35. Brown, C.J., Carrel, L., and Willard, H.F., (1997). Expression of genes from the human active and inactive X chromosomes. Am. J. Hum. Genet. 60, 1333–1343.

36. Carrel, L., and Willard, H.F. (1999). Heterogeneous gene expression from the inactive X chromosome: An X-linked gene that escapes X inactivation in some human cell lines but is inactivated in others. Proc. Natl. Acad. Sci. U. S. A. 96, 7364–7369.

37. Pintacuda, G., and Cerase, A. (2015). X Inactivation Lessons from Differentiating Mouse Embryonic Stem Cells. Stem Cell Rev. Reports 11, 699–705.

38. Patel, S., Bonora, G., Sahakyan, A., Kim, R., Chronis, C., Langerman, J., Fitz-Gibbon, S., Rubbi, L., Skelton, R.J.P., Ardehali, R., et al. (2017). Human Embryonic Stem Cells Do Not Change Their X Inactivation Status during Differentiation. Cell Rep. 18, 54–67.

39. Fan, G., and Tran, J. (2011). X chromosome inactivation in human and mouse pluripotent stem cells. Hum. Genet. 130, 217–222.

40. Sierra, I., and Anguera, M.C. (2019). Enjoy the silence: X-chromosome inactivation diversity in somatic cells. Curr. Opin. Genet. Dev. 55, 26–31.

41. Syrett, C.M., Paneru, B., Sandoval-Heglund, D., Wang, J., Banerjee, S., Sindhava, V., Behrens, E.M., Atchison, M., and Anguera, M.C. (2019). Altered X-chromosome inactivation in T cells may promote sex-biased autoimmune diseases. JCI Insight 4, e126751.

42. Beyer, A.I., and Muench, M.O. (2017). Comparison of human hematopoietic reconstitution in different strains of immunodeficient mice. Stem Cells Dev. 26, 102–112.

43. Yu, B., Qi, Y., Li, R., Shi, Q., Satpathy, A.T., and Chang, H.Y. (2021). B cell-specific XIST complex enforces X-inactivation and restrains atypical B cells. Cell 184, 1790–1803.e17.

44. Pereira, J.P., Girard, R., Chaby, R., Cumano, A., and Vieira, P. (2003). Monoallelic expression of the murine gene encoding Toll-like receptor 4. Nat. Immunol. 4, 464–470.

45. Gendrel, A.V., Marion-Poll, L., Katoh, K., and Heard, E. (2016). Random monoallelic expression of genes on autosomes: Parallels with X-chromosome inactivation. Semin. Cell Dev. Biol. 56, 100–110.

46. Mould, A.W., Pang, Z., Pakusch, M., Tonks, I.D., Stark, M., Carrie, D., Mukhopadhyay, P., Seidel, A., Ellis, J.J., Deakin, J., et al. (2013). Smchd1 regulates a subset of autosomal genes subject to monoallelic expression in addition to being critical for X inactivation. Epigenetics and Chromatin 6, 19.

47. Chow, J.C., Ciaudo, C., Fazzari, M.J., Mise, N., Servant, N., Glass, J.L., Attreed, M., Avner, P., Wutz, A., Barillot, E., et al. (2010). LINE-1 activity in facultative heterochromatin formation during X chromosome inactivation. Cell 141, 956–969.

48. Allen, E., Horvath, S., Tong, F., Kraft, P., Spiteri, E., Riggs, A.D., and Marahrens, Y. High concentrations of long interspersed nuclear element sequence distinguish monoallelically expressed genes.

49. Ayachi, S., Buscarlet, M., and Busque, L. (2020). 60 Years of clonal hematopoiesis research: From X-chromosome inactivation studies to the identification of driver mutations. Exp. Hematol. 83, 2–11.

50. Eldredge, N., and Gould, S.J. (1972). Punctuated Equilibria: An Alternative to Phyletic Gradualism. In Models in Paleobiology, pp. 82–115.

51. Castel, S.E., Levy-Moonshine, A., Mohammadi, P., Banks, E., and Lappalainen, T. (2015). Tools and best practices for data processing in allelic expression analysis. Genome Biol. 16.

52. Gupta, S., Lafontaine, D.L., Vigneau, S., Vinogradova, S., Mendelevich, A., Igarashi, K.J., Bortvin, A., Alves-Pereira, C.F., Clement, K., Pinello, L., et al. (2020). DNA methylation is a key mechanism for maintaining monoallelic expression on autosomes. bioRxiv, 10.1101/2020.02.20.954834.

